# Valid statistical approaches for clustered data: A Monte Carlo simulation study

**DOI:** 10.1101/2020.11.27.400945

**Authors:** Kristen A. McLaurin, Amanda J. Fairchild, Dexin Shi, Rosemarie M. Booze, Charles F. Mactutus

## Abstract

The translation of preclinical studies to human applications is associated with a high failure rate, which may be exacerbated by limited training in experimental design and statistical analysis. Nested experimental designs, which occur when data have a multilevel structure (e.g., *in vitro:* cells within a culture dish; *in vivo:* rats within a litter), often violate the independent observation assumption underlying many traditional statistical techniques. Although previous studies have empirically evaluated the analytic challenges associated with multilevel data, existing work has not focused on key parameters and design components typically observed in preclinical research. To address this knowledge gap, a Monte Carlo simulation study was conducted to systematically assess the effects of inappropriately modeling multilevel data via a fixed effects ANOVA in studies with sparse observations, no between group comparison within a single cluster, and interactive effects. Simulation results revealed a dramatic increase in the probability of type 1 error and relative bias of the standard error as the number of level-1 (e.g., cells; rats) units per cell increased in the fixed effects ANOVA; these effects were largely attenuated when the nesting was appropriately accounted for via a random effects ANOVA. Thus, failure to account for a nested experimental design may lead to reproducibility challenges and inaccurate conclusions. Appropriately accounting for multilevel data, however, may enhance statistical reliability, thereby leading to improvements in translatability. Valid analytic strategies are provided for a variety of design scenarios.

## Introduction

Preclinical studies, which range from molecular and *in vitro* studies to *in vivo* studies utilizing biological systems to model disease [1], are not immune [2–3] from the well-documented reproducibility issues observed in clinical fields [4]. Various factors, including rigorous standardization of preclinical experiments [e.g., 5–6], lack of scientific rigor [e.g., 7–8], and bias [e.g., Publication Bias: 9–10; Reporting Bias: 11], threaten reproducibility in preclinical science. Moreover, utilization of inappropriate statistical techniques is pervasive in the basic biological sciences [12–13]; a factor that likely exacerbates the reproducibility crisis.

Although statistical analyses have become an essential component of scientific publications [14], basic scientists receive limited training in experimental design and quantitative methodology [14–15]. When doctoral curriculums include training in statistics, introductory courses primarily focus on traditional quantitative techniques (e.g., analysis of variance; ANOVA), but often fail to cover specialized statistical methods that are integral to contemporary research [e.g., multilevel modeling; 16]. For example, clustered data (e.g., *in vitro*: cells within a culture dish; *in vivo*: rats within a litter; see Fig 1), which are prevalent in preclinical research [17–18], often violate the independent observation assumption underlying many traditional statistical techniques (e.g., *t*-tests, ANOVA). Multilevel modeling [also called hierarchical linear modeling; 19], however, appropriately accounts for the shared variance in nested data, thereby precluding violations of the independent observation assumption. For nearly fifty years [20–22], preclinical scientists have recognized the importance of appropriately defining the experimental unit, and yet a majority of preclinical studies continue to inappropriately analyze clustered data [e.g., Animal Models: 23–25, Developmental Psychobiology: 18, Neuroscience: 17].

**Fig 1.**
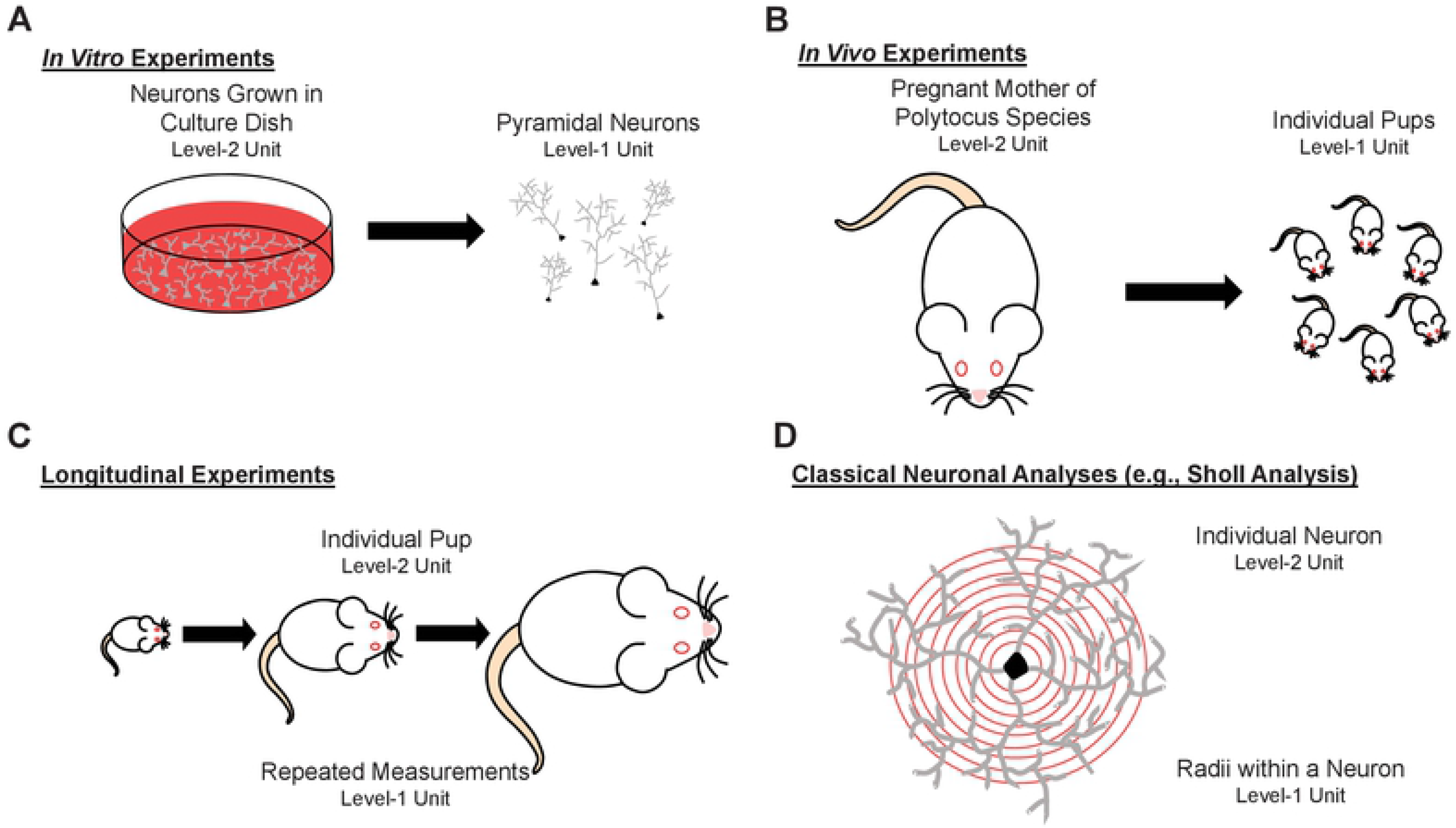
Examples of nested data commonly observed in preclinical studies. Nested data occurs when multiple subjects and/or measurements are obtained from a single higher-order group. Examples of nested data range from *in vitro* experiments (i.e., cells within a culture dish **(A)**) to *in vivo* experiments utilizing polytocus species (e.g., rat pups within a litter **(B)**). Multilevel data can also occur with the use of longitudinal experimental designs (i.e., repeated measurements are taken from a single individuals **(C)**) or the classical Sholl analysis (i.e., radii are nested within a neuron **(D)**; [62–63]).

Within preclinical fields, simulation studies have afforded an opportunity to empirically evaluate the implications of inappropriately modeling clustered data [e.g., 17–18, 26–28]. Spuriously significant effects, evidenced by inflated type 1 error rates [17–18, 26–28], are a well-recognized consequence of inappropriately modeling clustered data. With regards to statistical power, higher intraclass correlation coefficients (ICC; i.e., the relatedness of nested data [29–30]) are associated with lower statistical power [17]. To date, however, the majority of work examining statistical implications of violating the independent observation assumption has primarily considered parameters and design components not well aligned with those observed in preclinical studies (i.e., overly large sample and cluster size), as well as simplified models that preclude the examination of interactive effects. Moreover, the effect of inappropriately modeling multilevel data on the statistical accuracy of parameter estimates has not yet been systematically evaluated under these conditions.

In light of gaps in previous work, a Monte Carlo simulation study was conducted to empirically evaluate the effects of inappropriately modeling multilevel data using parameters more reflective of preclinical work. Specifically, the study considered: 1) sparse data, defined by either a small number of level-1 units per cell [31] or a small number of clusters; 2) no between group comparison within clusters, and 3) interactive effects. The rationale of including the latter derives from requirements by the National Institutes of Health to include sex as a biological variable (NOT-OD-15-102). Population data in line with a fully-crossed two-factor ANOVA, where treatment units were nested within clusters, was simulated to consider the impact on both main effects (e.g., treatment and sex) and interaction terms (e.g., treatment x sex). Study outcomes were compared across a traditionally-used fixed effects ANOVA model and a two-level random effects ANOVA model that allowed variation in both the intercept and slope. Outcome variables were selected to assess both the accuracy of hypothesis testing and parameter estimates in the model. Valid analytic strategies are provided for a variety of design scenarios. Given the current rigor and reproducibility crisis in the biomedical sciences, evaluating the implications of inappropriate statistical practices is integral to the quest for more efficient and reliable data.

## Results

The population model in the simulation was a fully crossed 2×2 random effects ANOVA model, with two binary predictors and an interaction term. Population parameters for level-1 sample size (i.e., number of level-1 units per cell; *N*), level-2 sample size (i.e., number of clusters; *C*), the parameter effect size for the main effect (β_1_), the parameter effect size for the interaction effect (β_3_), and ICC were systematically varied yielding a 6 x 5 x 4 x 4 x 2 factorial design with 960 conditions (Table 1). Each condition was replicated 1,000 times, yielding 960,000 datasets for analysis. Given the extremely large sample size, and corresponding inflation of statistical significance, practical significance was evaluated against a partial η^2^ ≥ 0.01 criterion, indicating that at least 1% of the variance in a given outcome was attributable to the effect of interest [32]. The partial η^2^ values for each parameter, and all possible interactions among the parameters, are presented for all outcome variables in Table 2 (Main Effect of β_1_) and Table 3 (Interaction Effect of β_3_).

**Table 1.** Simulation Population Parameters and Corresponding Levels of Each Parameter.

| Population Parameter | Levels |
| --- | --- |
| Sample Size: Level-1 ($N$) | 2, 4, 6, 8, 10, 12 |
| Sample Size: Level-2 ($C$) | 4, 8, 12, 16, 20 |
| Parameter Effect Size |  |
| Main Effect of $\beta_1$ ($\beta_1$) | 0, 0.14, 0.39, 0.59 |
| Interaction Effect $\beta_3$ ($\beta_3$) | 0, 0.14, 0.39, 0.59 |
| Intraclass Correlation (ICC) | 0.16, 0.6 |

**Table 2.**
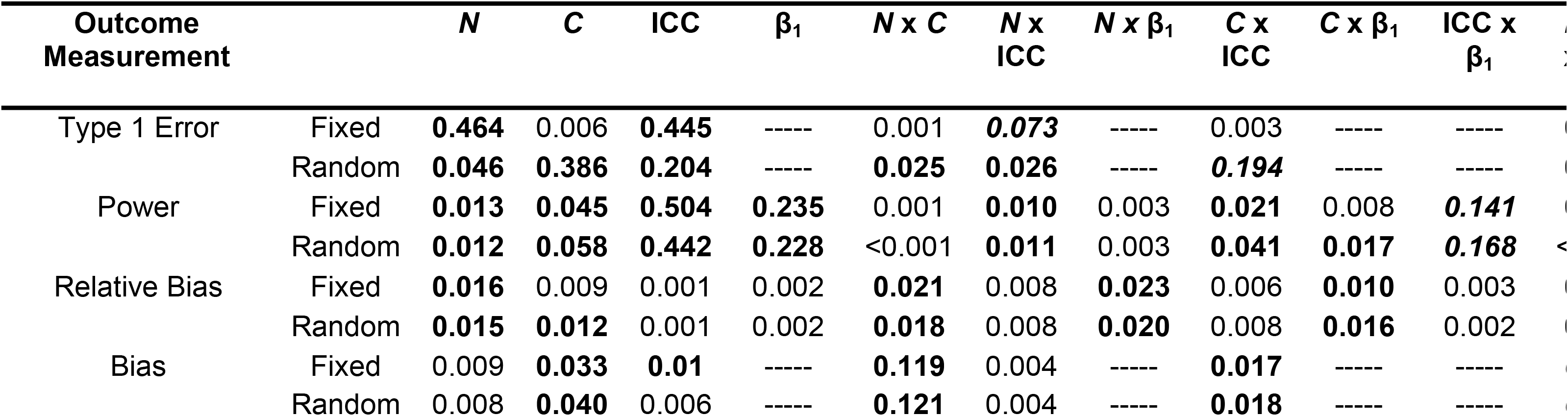
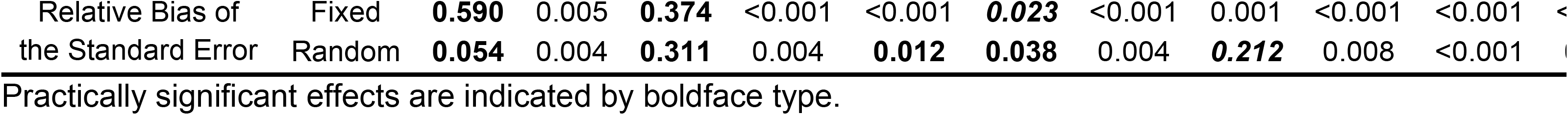
Partial η^2^ for the Main Effect of β_1_.

**Table 3.** Partial η^2^ for the Main Effect of β_3_.

| Outcome<br>Measurement | | *N* | *C* | ICC | $\beta_3$ | *N* x *C* | *N* x<br>ICC | *N* x $\beta_3$ | *C* x<br>ICC | *C* x $\beta_3$ |
| --- | --- | --- | --- | --- | --- | --- | --- | --- | --- | --- |
| Type 1 Error | Fixed | **0.602** | **0.040** | **0.109** | ---- | 0.008 | **0.197** | ---- | **0.018** | ---- |
|  | Random | **0.027** | **0.105** | **0.555** | ---- | **0.013** | 0.004 | ---- | **0.203** | ---- |
| Power | Fixed | **0.067** | **0.064** | **0.309** | **0.233** | 0.006 | **0.038** | **0.028** | **0.040** | **0.028** |
|  | Random | **0.042** | **0.082** | **0.285** | **0.235** | 0.006 | **0.039** | **0.019** | **0.054** | **0.038** |
| Relative Bias | Fixed | 0.004 | 0.004 | <0.001 | <0.001 | **0.014** | 0.007 | **0.020** | 0.004 | 0.004 |
|  | Random | 0.003 | 0.003 | <0.001 | <0.001 | **0.020** | 0.006 | **0.018** | 0.003 | 0.003 |
| Bias | Fixed | **0.023** | **0.013** | 0.004 | ---- | **0.077** | **0.013** | ---- | **0.019** | ---- |
|  | Random | **0.018** | **0.023** | 0.004 | ---- | **0.077** | **0.014** | ---- | **0.019** | ---- |
| Relative Bias of<br>the Standard Error | Fixed | **0.684** | **0.012** | 0.002 | <0.001 | <0.001 | **0.286** | <0.001 | 0.005 | <0.001 |
|  | Random | **0.026** | **0.050** | **0.577** | 0.002 | 0.005 | **0.011** | 0.003 | **0.187** | 0.003 |

Practically significant effects are indicated by boldface type.

### Type 1 error

For experimental conditions where the population value of interest (i.e., β_1_, β_3_) was zero, the accuracy of hypothesis testing was evaluated using type 1 error, which was defined as the proportion of replications in a given condition that yielded statistically significant results. Type 1 error was evaluated against a nominal α criterion of 0.05.

#### Main effect (β_1_)

For the main effect of β_1_ in the fixed effects model (Fig 2A), the probability of type 1 error ranged from 5.8% to 23.3% for the small ICC and from 10.3% to 49.8% for the large ICC. The probability of type 1 error rates increased as the number of level-1 units per cell increased, but the rate of increase was dependent upon ICC [*N* x ICC Interaction: η 2=0.073]. Specifically, the probability of type 1 error increased at a significantly greater rate when the ICC was large relative to a small ICC [First Order Polynomial: *R*^2^s>0.91; *F*(1,236)=351.1, *p*≤0.001]. Most critically, however, observed type 1 error rates were greater than the established α criterion of 0.05 across all levels of the population parameters (i.e., *N* and ICC).

**Fig 2.**
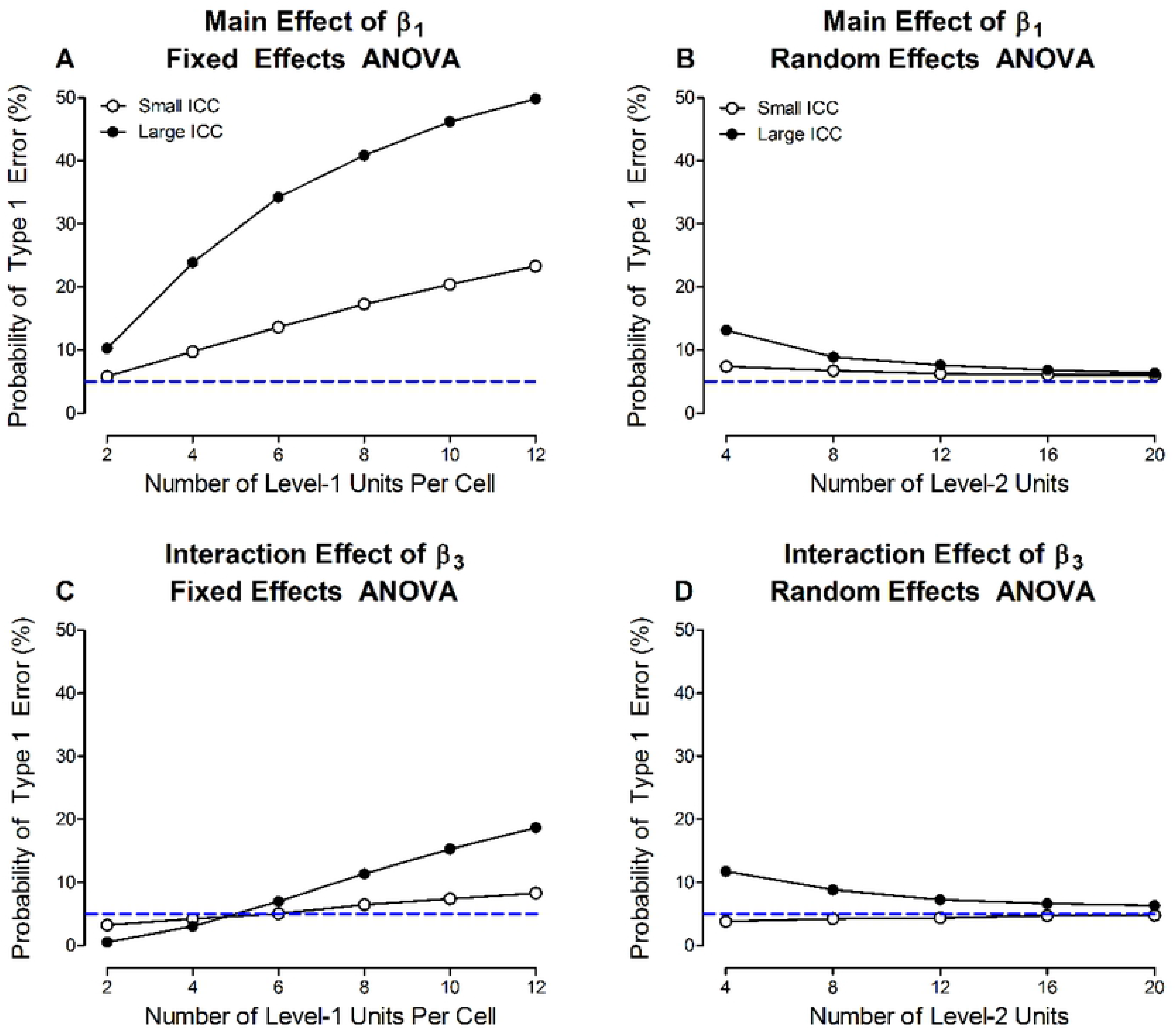
Probability of type 1 error. The probability of type 1 error (%) is illustrated as a function of β coefficient (i.e., Main Effect of β_1_: **A, B**; Interaction Effect of β_3_: **C, D**), analytic approach (i.e., Fixed Effects ANOVA: **A, C**; Random Effects ANOVA: **B, D**), and intraclass correlation (ICC). In the fixed effects ANOVA **(A,C**), mean estimates for the probability of type 1 error increased as the number of level-1 units per cell increased; estimates which were greater than the established α criterion of 0.05. Utilization of a random effects ANOVA **(B,D**), however, improved the accuracy of hypothesis testing evidenced by type 1 error rates that approximate the established α criterion. The dashed blue line reflects the established α criterion of 0.05.

When the nested experimental design was appropriately accounted for via a random effects model, elevated type 1 error rates were largely attenuated (Fig 2B). In the random effects model, mean type 1 error rates ranged from 6% to 7.3% for the small ICC and from 6.3% to 13.2% for the large ICC; these values were dependent upon an interaction between the number of level-2 units and ICC [*C* x ICC Interaction: η 2=0.194]. Specifically, the probability of type 1 error decreased at a significantly faster rate when the ICC was large relative to a small ICC [First Order Polynomial: *R*^2^s>0.83; *F*(1, 236)=112, *p*≤0.001].

#### Interaction effect (β_3_)

With regard to the interaction effect of β_3_ in the fixed effects model (Fig 2C), mean estimates for type 1 error ranged from 3.3% to 8.3% for the small ICC and from 0.5% to 18.7% for the large ICC. Consistent with observations for β_1_, mean estimates for the probability of type 1 error were dependent upon an interaction between the number of level-1 units per cell and the value of the ICC [*N* x ICC Interaction: η 2=0.197]. Specifically, as the number of level-1 units per cell increased, the probability of type 1 error increased; an increase that was significantly faster when the ICC was large relative to a small ICC [First Order Polynomial: *R*^2^s>0.99; *F*(1, 236)=490.7, *p*≤0.001]. Type 1 error rates were conservative when there were only two level-1 units per cell. There was diminished accuracy of hypothesis testing when more than four level-1 units per cell were selected, however, evidenced by a type 1 error rate that was greater than the established α criterion of 0.05.

Utilization of a random effects model, to appropriately account for the nested experimental design, largely attenuated the elevated type 1 error rates for the interaction effect of β_3_ (Fig 2D). When the ICC was small, mean estimates for the probability of type 1 error in the random effects model ranged from 3.8% to 4.8%; observations which support accurate estimates across all level-2 population parameters. For the large ICC, mean estimates for the probability of type 1 error in the random effects model ranged from 6.3% to 11.7%; observations which revealed a greater probability of type 1 error when fewer level-2 units per cell were sampled. The overall ANOVA confirmed our observations, revealing a practically significant interaction between the number of level-2 units and the ICC [*C* x ICC Interaction: η 2=0.203].

### Power

For experimental conditions where the population value of interest (i.e., β_1_, β_3_) was non-zero, the accuracy of hypothesis testing was assessed via statistical power, which was defined by the proportion of replications in a given condition that yielded statistically significant results. Statistical power was evaluated against a criterion of 0.80 [32].

#### Main effect (β_1_)

With regard to the main effect of β_1_ in the fixed effects model (Fig 3A), statistical power ranged from 12.2% to 73.8% for the small ICC and from 0.03% to 8.8% for the large ICC. A practically significant interaction between the magnitude of β_1_ and ICC was observed [β_1_ x ICC Interaction: η_p_^2^=0.141]. As the magnitude of β_1_ increased, statistical power increased; an increase that was significantly faster when the ICC was small relative to a large ICC [First Order Polynomial: *R*^2^s>0.97; *F*(1,716)=734.5, *p*≤0.001]. However, the observed statistical power failed to reach the established criterion of 0.80 at any levels of the population parameters studied.

**Fig 3.**
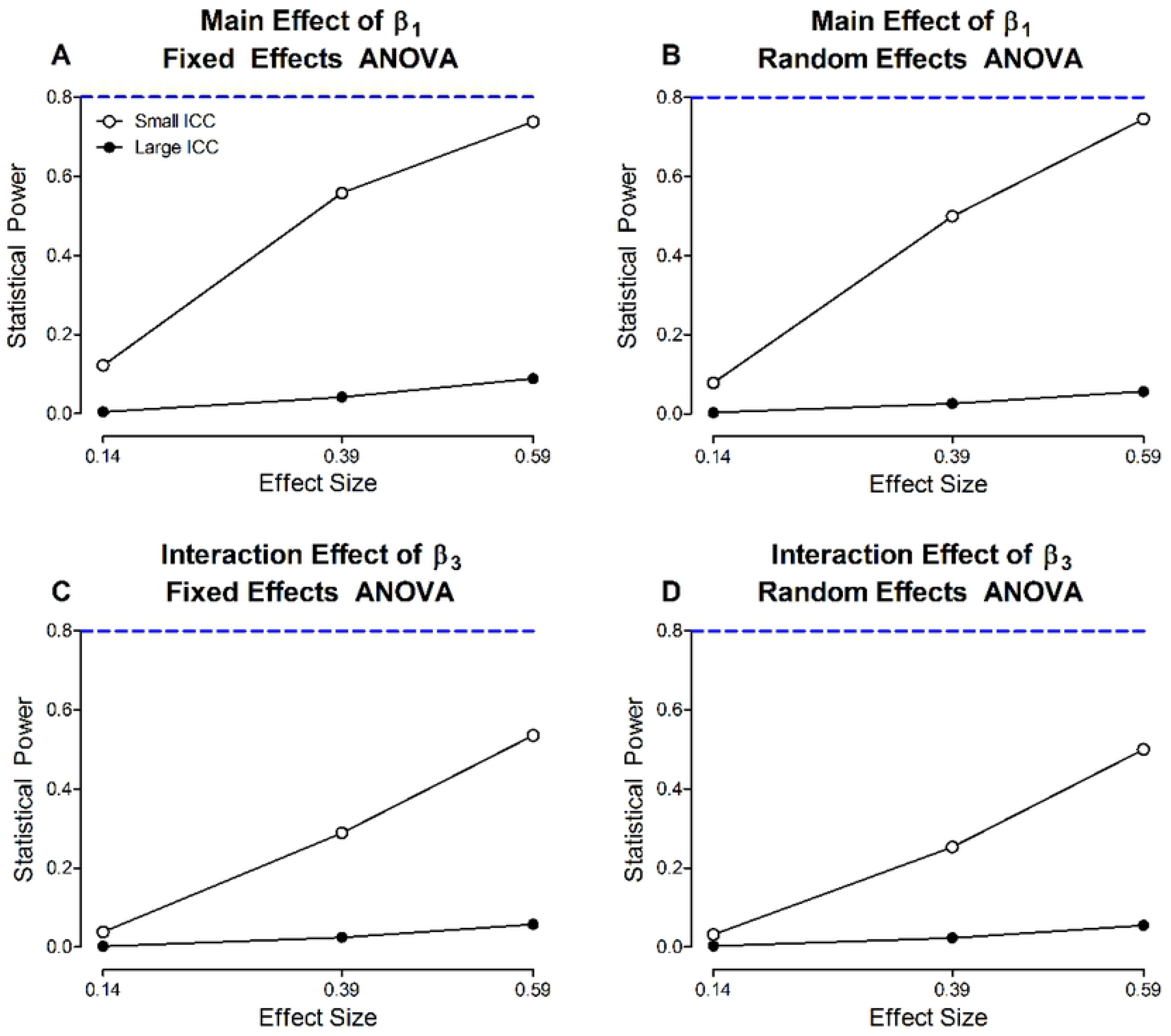
Statistical power. Statistical power is illustrated as a function of coefficient magnitude (i.e., 0.14, 0.39, 0.59), β coefficient (i.e., Main Effect of β_1_: **A, B**; Interaction Effect of β_3_: **C, D**), analytic approach (i.e., Fixed Effects ANOVA: **A,C**; Random Effects ANOVA: **B,D**), and intraclass correlation (ICC). Independent of analytic approach and/or β coefficient, statistical power failed to reach the established criterion of 0.80. Overall, statistical power was lower for the interaction effect of β_3_. The dashed blue line reflects the established criterion of 0.80.

Utilizing a random effects model to appropriately account for the nested experimental design did not significantly improve the statistical power to detect effects. In the random effects model, statistical power ranged from 7.8% to 74.6% for the small ICC and from 0.03% to 5.6% for the large ICC (Fig 3B); these estimates were dependent upon an interaction between the magnitude of β_1_ and ICC [β_1_ x ICC Interaction: η 2=0.168]. Consistent with observations for the fixed effects ANOVA, statistical power to detect the main effect of β_1_ increased at a significantly faster rate when the ICC was small relative to a large ICC [First Order Polynomial: *R*^2^s>0.97; *F*(1, 716)=720.2, *p*≤0.001]. Although statistical power failed to reach the established criterion of 0.80 in the random effects model, it is noteworthy that the utilization of an appropriate, advanced quantitative method had no adverse effects (i.e., did not decrease) on statistical power.

#### Interaction effect (β_3_)

For the interaction effect in the fixed effects model (Fig 3C), statistical power ranged from 3.8% to 53.6% for the small ICC and from 0% to 5.7% for the large ICC. These estimates were dependent upon an interaction between the magnitude of β_3_ and the ICC [β_3_ x ICC Interaction: η_p_^2^=0.150] and were lower for the interaction effect of β_3_ relative to the main effect of β_1_. As the magnitude of β_3_ increased, statistical power increased; an increase that was significantly faster when the ICC was small relative to a large ICC [First Order Polynomial: *R*^2^s>0.97; *F*(1, 716)=345.2, *p*≤0.001].

Utilization of a random effects model to appropriately account for the nested experimental design did not increase statistical power to detect effects. In the random effects model, statistical power ranged from 3.1% to 50% for the small ICC and from 0% to 5.5% for the large ICC (Fig 3D). Consistent with observations for the interaction effect of β_3_ in the fixed effects model, statistical power was dependent upon an interaction between the magnitude of β_3_ and ICC [β_3_ x ICC Interaction: η 2=0.150]. Statistical power increased at a significantly faster rate when the ICC was small relative to a large ICC [First Order Polynomial: *R*^2^s>0.96; *F*(1, 716)=318.5, *p*≤0.001]. Consistent with observations for β_1_, statistical power for the interaction effect of β_3_ failed to reach the established criterion of 0.80 in either the fixed effects model or the random effects model; results that suggest preclinical studies may often be underpowered, resulting in decreased accuracy for hypothesis testing.

### Relative bias of parameter estimates

For conditions where the parameter effect size of interest (i.e., β_1_, β_3_) was non-zero, the relative bias of parameter estimates was evaluated to assess the accuracy of model parameter estimates. Relative bias was defined as the difference between the observed sample estimate and the true value of a given parameter, relative to the true value of the parameter being estimated:

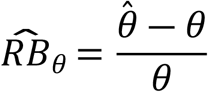

where 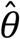 is the average parameter estimate across 1000 replications and θ refers to the population parameter value. Values of relative bias that exceeded |10%| were considered poor [33].

#### Main effect (β_1_)

For the main effect of β_1_, a practically significant four-way interaction between the number of level-1 units per cell, the number of level-2 units, the magnitude of β_1_, and ICC [*N* x *C* x β_1_ x ICC Interaction: η_p_^2^=0.043] was observed in the fixed effects model. Under conditions where the ICC was small, results demonstrated tolerable rates of relative bias across varying values of β_1_, with mean estimates for relative bias ranging from −6.6% to 7.2% when β_1_=0.14, from −5.7% to 1.6% when β_1_=0.39, and from −1.7% to 2.1% when β_1_=0.59. For the large ICC (Fig 4A, 4C, 4E), however, excessive rates of bias were observed under conditions when the magnitude of β_1_ was small. Specifically, mean estimates for relative bias ranged from −21.2% to 38% when β_1_=0.14. This contrasted to relative bias estimates in the large ICC conditions when β_1_ was either medium (0.39) or large (0.59), with relative bias estimates ranging from −5.8% to 8.7% and from −6.5% to 6.9%, respectively. Overall, the pattern of relative bias was random, centered around zero, and did not consistently exceed the established criterion of |10%| in these conditions.

**Fig 4.**
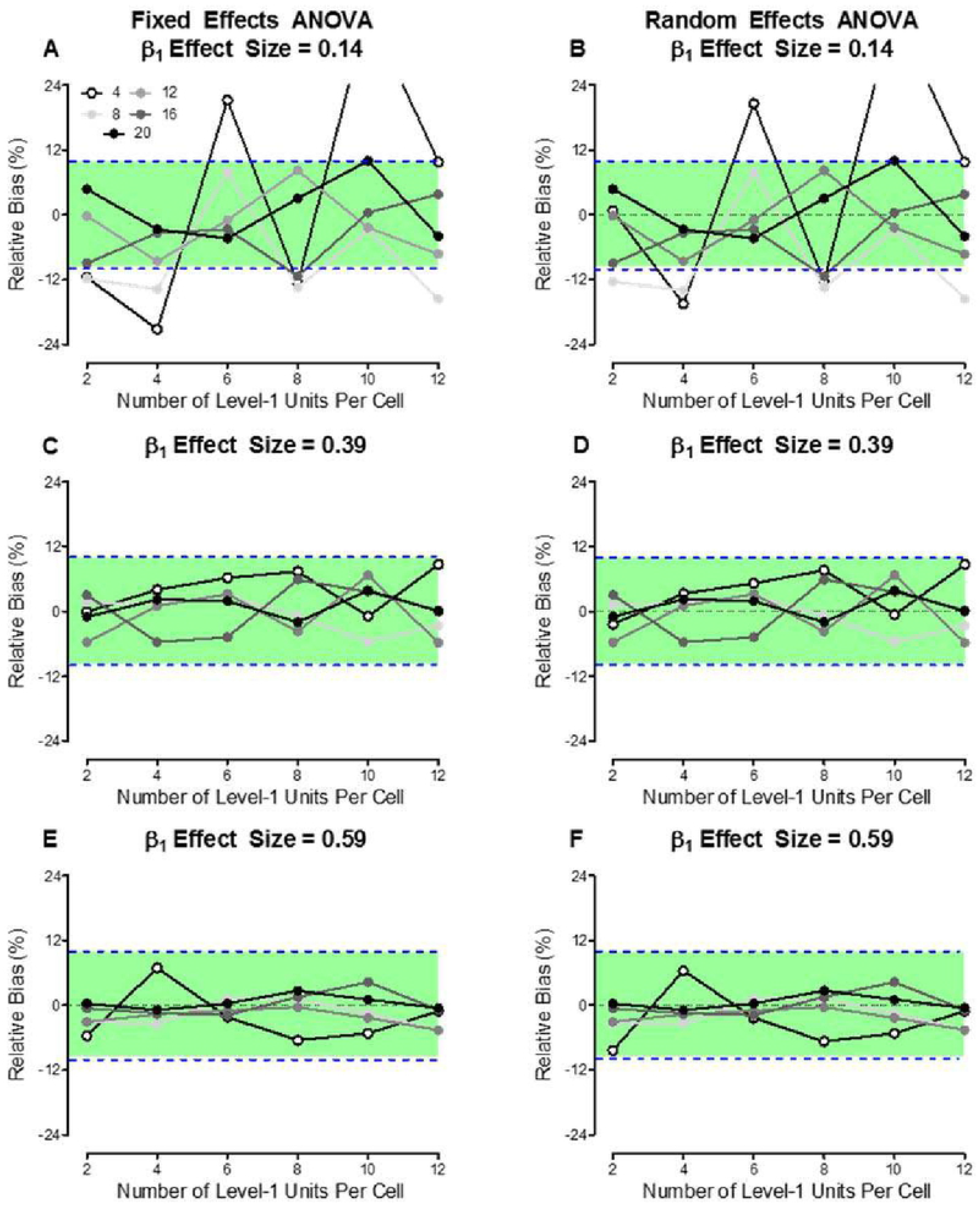
Relative bias of parameter estimates. Relative bias (%) is illustrated for the main effect of β_1_ and the large intraclass correlation as a function of coefficient magnitude (i.e., 0.14 (**A, B**), 0.39 (**C, D**), 0.59 (**E, F**)), analytic approach (i.e., Fixed Effects ANOVA: **A, C, E**; Random Effects ANOVA: **B, D, F**), the number of level-1 units per cell, and the number of level-2 units. Independent of analytic approach, elevated rates of relative bias were observed for some conditions when the magnitude of β_1_ was small (**A,B**). Overall, however, the pattern of relative bias was random, centered around zero, and not consistently exceed the established criterion of |10%| for either the fixed effects or random effects ANOVA. The green area within the two dashed blue lines reflects the acceptable levels of relative bias. Points outside of the green area are greater than the established criterion of |10%|.

With regard to the main effect of β_1_ in the random effects model, a practically significant four-way interaction between the number of level-1 units per cell, the number of level-2 units, the magnitude of β_1_, and ICC [*N* x *C* x β_1_ x ICC Interaction: η_p_^2^=0.041] was also observed. For the small ICC, results demonstrated tolerable rates of relative bias across values of β_1_, with mean estimates for relative bias ranging from −6.6% to 7.3% when β_1_=0.14 from −3.1% to 1.9% when β_1_=0.39, and from −1.4% to 2.2% when β_1_=0.59. Under parameter conditions where the ICC was large (Fig 4B, 4D, 4F), results demonstrated intolerable relative bias when the magnitude of β_1_ was small, with estimates ranging from −16.5% to 37.9%. This contrasted to relative bias estimates when the magnitude of β_1_ was either medium (0.39) or large (0.59) in these conditions, with relative bias estimates ranging from −5.7% to 8.7% and from −8.4% to 2.7%, respectively. The comparability of relative bias results across the random effects and fixed effects models indicate neither a beneficial nor detrimental effect of appropriately modeling nested data on the relative bias of model parameter estimates.

#### Interaction effect (β_3_)

For the interaction effect of β_3_, a practically significant four-way interaction between the number of level-1 units per cell, the number of level-2 units, the magnitude of β_3_, and ICC [*N* x *C* x β_3_ x ICC Interaction: η_p_^2^=0.033] was observed in the fixed effects model. When β_3_=0.14, mean estimates for relative bias ranged from −17.8% to 11.1% and from −23.4% to 21.3% for the small ICC and large ICC, respectively. When β_3_=0.39, mean estimates for relative bias ranged from −2.0% to 1.9% for the small ICC and from −8.4% to 8.6% for the large ICC. Finally, when β_3_=0.59, mean estimates for relative bias ranged from −1.8% to 1.3% and from −5.4% to 11.7% for the small ICC and large ICC, respectively. Overall, relative bias did not consistently exceed the established criterion of |10%|. Notably, however, there were excessive rates of relative bias when the magnitude of the interaction term was small.

Mean estimates for relative bias for the interaction effect of β_3_ in the random effects model approximated those observed in the fixed effects model. Furthermore, consistent with observations for the interaction effect of β_3_ in the fixed effects ANOVA, a practically significant interaction between the number of level-1 units per cell, the number of level-2 units, the magnitude of β_3_, and the ICC was observed [*N* x *C* x β_3_ x ICC Interaction: η_p_^2^=0.030]. Specifically, for the small ICC, mean estimates for relative bias ranged from −23.9% to 10.6% when β_3_=0.14, from −1.8% to 2.6% when β_3_=0.39, and from −1.7% to 1.9% when β_3_=0.59. Under parameter conditions where the ICC was large, mean estimates for relative bias ranged from −23.4% to 20.5% when β_3_=0.14, from −8.3% to 9.8% whenβ_3_=0.39, and from −5.7% to 11.6% when β_3_=0.59. As with results for the fixed effects model, there was a modest elevation of relative bias when the magnitude of the interaction term was small. Overall however, and consistent with observations for β_1_, relative bias for the interaction effect of β_3_ did not consistently exceed the established criterion of |10%| in either the fixed effects model or the random effects model.

### Absolute bias of parameter estimates

When relative bias was undefined in experimental conditions (i.e., when true values of the parameter effect size of the β coefficients were equal to zero), absolute bias was calculated as follows:

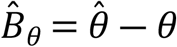

Values of absolute bias that exceeded |10%| were considered poor [33].

#### Main Effect (β_1_)

For the main effect of β_1_ in the fixed effects model, mean estimates for absolute bias ranged from −0.8% to 1.3% for the small ICC and from −2.5% to 6.2% for the large ICC. A practically significant interaction between the number of level-1 units per cell, the number of level-2 units, and ICC was observed [*N* x *C* x ICC Interaction: η_p_^2^=0.060].

Utilizing a random effects model to appropriately account for the nested experimental design had neither a beneficial nor adverse effect on absolute bias. Consistent with observations for the main effect of β_1_ in the fixed effects ANOVA, a practically significant interaction between the number of level-1 units per cell, the number of level-2 units and ICC was observed [*N* x *C* x ICC Interaction: η_p_^2^=0.058]. Specifically, for the small ICC, mean estimates for absolute bias ranged from −0.8% to 1.5%. For the large ICC, mean estimates for absolute bias ranged from −2.0% to 6.4% in the random effects ANOVA. Therefore, absolute bias did not exceed established criterion of |10%| for any conditions assessed in either the fixed effects ANOVA or the random effects ANOVA.

#### Interaction Effect (β_3_)

With regards to the interaction effect of β_3_, a practically significant three-way interaction between the number of level-1 units per cell, the number of level-2 units, and ICC [*N* x *C* x ICC Interaction: η_p_^2^=0.028] was observed in the fixed effects model. Mean estimates for absolute bias ranged from −1.2% to 1.1% and from −2.3% to 1.8% for the small ICC and large ICC, respectively.

Utilizing a random effects model to appropriately account for the nested experimental design had neither a beneficial nor adverse effect on absolute bias. For the small ICC, mean estimates for absolute bias ranged from −1.1% to 1.1%. For the large ICC, mean estimates for absolute bias ranged from −3.7% to 1.1%. Consistent with observations for the main effect of β_1_, absolute bias for the interaction effect of β_3_ did not exceed the established criterion of |10%| for any conditions assessed in either the fixed effects ANOVA or the random effects ANOVA.

### Relative bias of the standard error

Relative bias of the standard error was evaluated for all experimental conditions to examine the accuracy of error estimates using the following formula [34]:

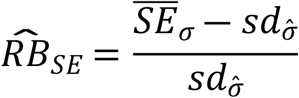

where 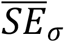 is the average standard error across replications and 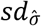 is the empirical standard deviation of parameter estimates. Values of relative bias of the standard error that were greater than |5%| were considered poor [35].

#### Main Effect (β_1_)

For the main effect of β_1_ in the fixed effects model (Fig 5A), results demonstrated intolerable levels of relative bias of the standard error across parameter combinations, ranging from −39.5% to −4.3% for the small ICC and from −65.2% to −14.8% for the large ICC. Mean estimates for relative bias of the standard error were dependent upon an interaction between the number of level-1 units per cell and ICC [*N* x ICC Interaction: η_p_^2^=0.023]. A one-phase decay provided a well-described fit for the relative bias of the standard error, independent of ICC (Small ICC: *R*^2^>0.99; Large ICC: *R*^2^>0.99). However, significant differences in the y-intercept [*F*(1,954)=156.1, *p*≤0.001], rate constant [i.e., K; *F*(1,954)=284.7, *p*≤0.001] and plateau [*F*(1,954)=106.4, *p*≤0.001] were observed. When failing to account for the nested data structure, the standard error for the main effect of β_1_ was negatively biased.

**Fig 5.**
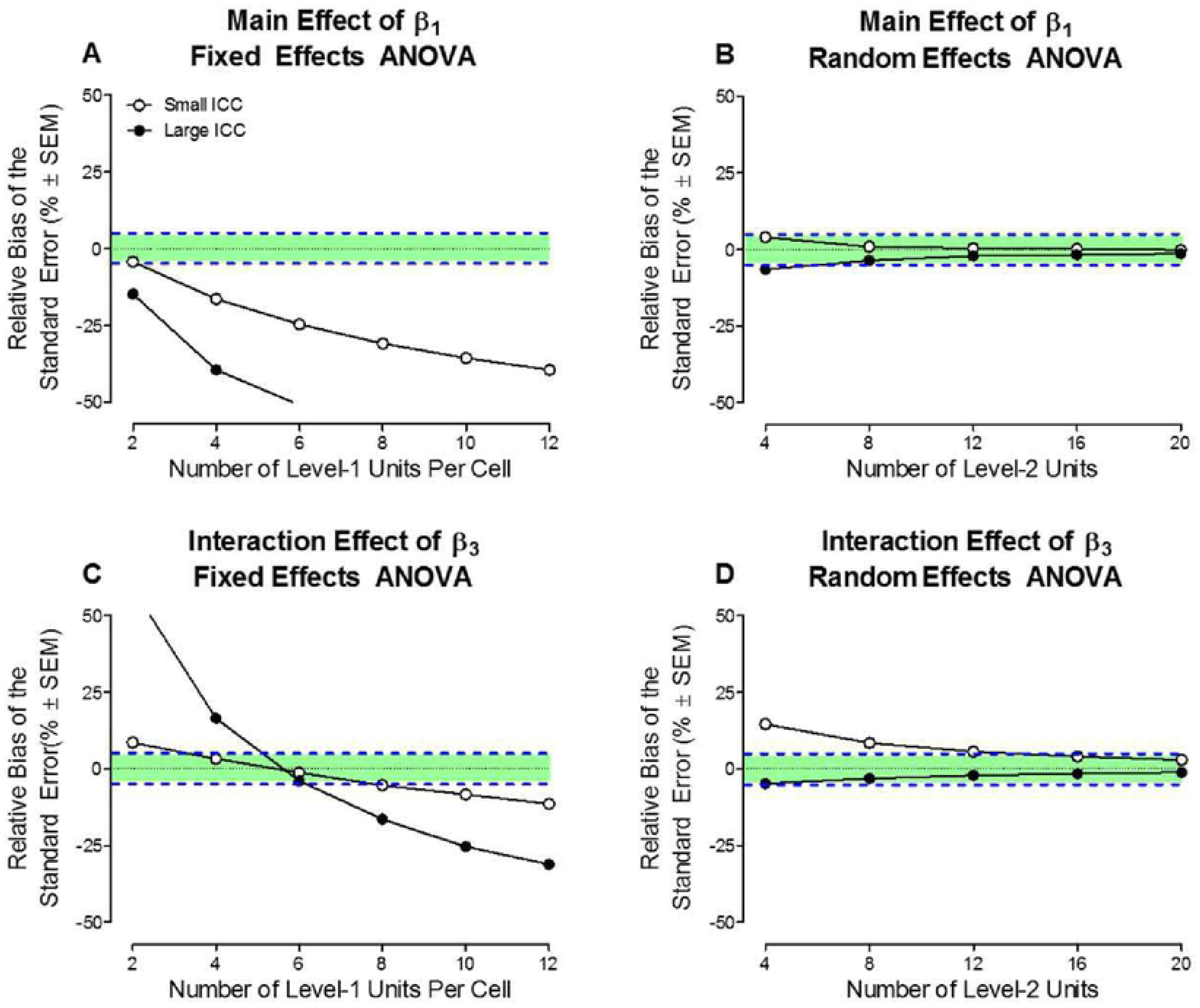
Relative bias of the standard error. Relative bias of the standard error (%) is illustrated as a function of β coefficient (i.e., Main Effect of β_1_: **A, B**; Interaction Effect of β_3_: **C, D**), analytic approach (i.e., Fixed Effects ANOVA: **A, C**; Random Effects ANOVA: **B, D**), and intraclass correlation (ICC). In the fixed effects ANOVA **(A,C)**, mean estimates for the relative bias of the standard error decreased as the number of level-1 units per cell increased supporting negatively biased standard errors. Utilization of a random effects ANOVA **(B,D)**, however, largely attenuated the relative bias of the standard error; an effect which represents a disattenuation of the standard error. The green area within the two dashed blue lines reflects the acceptable levels of relative bias of the standard error. Points outside of the green area are greater than the established criterion of |5%|.

When the nested experimental design was appropriately accounted for via a random effects model, however, the relative bias of the standard error was largely attenuated. When the ICC was small, mean estimates for the relative bias of the standard error in the random effects model ranged from 4% to −0.2%; estimates that were less than the established criterion of 5% across all level-2 units per cell. For the large ICC, mean estimates for the relative bias of the standard error ranged from −6.6% to −1.4%; observations which revealed a greater likelihood of biased standard errors when fewer level-2 units were sampled (Fig 5B). The overall ANOVA confirmed our observations, revealing a practically significant interaction between the number of level-2 units and the ICC [*C* x ICC Interaction: η_p_^2^=0.212]. Furthermore, an investigation of the empirical standard deviation of parameter estimates demonstrated negligible differences between the fixed effect models and random effect models across conditions. Thus, in line with previous methodological work, results demonstrated that utilization of a random effects model largely attenuated the relative bias of the standard error to approximate the established criterion of |5%|; an effect resulting from the disattenuation of the standard error.

#### Interaction Effect (β_3_)

For the interaction effect of β_3_ in the fixed effects model (Fig 5C), mean estimates for the relative bias of the standard error ranged from 8.6% to −11.4% for the small ICC and from 58.9% to −31.1% for the large ICC. Overall, the relative bias of the standard error was greater when the ICC was large relative to a small ICC. A shift in the direction of the relative bias of the standard error (i.e., from positively biased to negatively biased) was observed as the number of level-1 units per cell increased, in line with increased violations of independence. A practically significant interaction between the number of level-1 units per cell and ICC confirmed our observations [*N* x ICC Interaction: η_p_^2^=0.286]. A one-phase decay provided a well-described fit for the relative bias the standard error, independent of ICC (Small ICC: *R*^2^>0.99; Large ICC: *R*^2^>0.99). However, significant differences in the y-intercept [*F*(1,954)=599.2, *p*≤0.001], and rate constant [i.e., K; *F*(1,954)=93.0, *p*≤0.001] were observed. Consistent with the observations for β_1_ in the fixed effects model, when two or more level-1 units per cell were selected, there was diminished accuracy of standard error estimates.

Utilization of a random effects model to appropriately account for the nested experimental design, however, largely attenuated the relative bias of the standard error. In the random effects model, mean estimates for the relative bias of the standard error ranged from 14.6% to 3% for the small ICC and from −4.8% to −1.1% for the large ICC (Fig 5D). Independent of ICC, as the number of level-2 units increased, the relative bias of the standard error approached 0. The practically significant interaction between the number of level-2 units and the ICC [*C* x ICC Interaction: η_p_^2^=0.187] captures differences in the direction (i.e., Small ICC: positively biased; Large ICC: negatively biased) of relative bias of the standard error. Therefore, consistent with observations for β_1_, utilization of a random effects model largely disattenuated the standard error and had a negligible effect on the empirical standard deviation of parameter estimates; observations which support the implementation of random effects models when nested data are present in a design.

## Discussion

Inappropriately modeling clustered data via a fixed effects ANOVA promoted inaccurate hypothesis testing and artificially attenuated standard error estimates; both of these effects were largely mitigated when the nested data structure was appropriately accounted for via a random effects ANOVA. Spuriously significant effects, evidenced by type 1 error rates greater than the established α criterion of 0.05, were observed in the fixed effects ANOVA. Significant negatively biased standard errors, which artificially decrease estimates of the standard error, promoted inaccurate hypothesis testing in the fixed effects ANOVA. Notably, inappropriately modeling nested data had adverse effects on both the main effect of β_1_ and the interaction effect of β_3_; albeit the magnitude of these effects was dependent upon the β coefficient (i.e., β_1_ or β_3_) and outcome variable of interest. In contrast, appropriately modeling nested data via a random effects ANOVA improved the accuracy of both hypothesis testing (i.e., Type 1 Error) and parameter estimates (i.e., Relative Bias of the Standard Error). Statistical power failed to reach the established criterion of 0.8 in either the fixed effects or random effects ANOVA; a result reflecting the small sample sizes commonly utilized in preclinical research. Thus, failure to account for a nested experimental design has critical implications on inferential statistics and may hinder reproducibility in the behavioral and biomedical sciences.

Selection of two or more level-1 units per cell has prominent adverse effects on inferential statistics when analytic techniques fail to account for the nested data structure. Consistent with previous methodological work [e.g., 17–18, 26–28, 36–38], type 1 error rates were greater than the established α criterion of 0.05 in the fixed effects ANOVA; results which demonstrate that findings based on larger samples, different design characterizations, and simpler models (i.e., *t-*tests) extend to the types of parameters more commonly seen in preclinical studies. Notably, the profound negative bias in the standard error, which occurs even when the number of level-1 units per cell is small, likely promotes elevated type 1 error rates in the fixed effects ANOVA by decreasing within-group variance. When multilevel data is appropriately modeled via a random effects ANOVA, however, the type 1 error rate and relative bias of the standard error approximate the established criterion (i.e., α < 0.05 and |5%|, respectively).

Low statistical power has been recognized as a critical, albeit not universal, issue in preclinical research [39–40]. In the present simulation, statistical power failed to reach the established criterion of 0.8 in either the fixed effects or random effects ANOVA; a result reflecting the small level-1 and level-2 sample sizes modeled to reflect those commonly observed in preclinical studies [41–42]. To maximize statistical power in a nested experimental design, methodologists recommend increasing the number of level-2 units, rather than the number of level-1 units per cell [e.g., 28, 43]. However, given feasibility issues (e.g., time, cost) with increasing sample size, it is important to consider utilizing alternative experimental design strategies, including repeated-measures [44], the inclusion of covariates [45–47], and use of no dependent observations [18], to increase statistical power. Implementation of these strategies is especially important in light of requirements by the NIH to include sex as a biological variable (NOT-OD-15-102); a requirement that necessitates investigation of interaction terms, which exhibit lower statistical power than main effects.

The assessment of two ICC variants revealed the importance of the value of ICC across all outcome measures. Specifically, in the fixed effects ANOVA, the value of ICC altered the magnitude, but not the presence, of inaccurate hypothesis testing and parameter estimates. The importance of calculating and reporting the ICC in preclinical studies, therefore, cannot be understated. ICC (i.e., ρ; [29–30]), which reflects the relatedness of nested data, is calculated by dividing the between-cluster variability by the total variability (i.e., within-cluster variability and between-cluster variability; [19]). Values of ICC range from zero to one, whereby, a higher ICC represents increased similarity within a cluster. Given that even small ICC values (i.e., ρ < 0.05) may have critical implications on inferential statistics [48–49], researchers should also conduct a formal statistical test to determine whether the ICC is statistically significant. Winer [50] and Denenberg 51] proposed a preliminary test to calculate an *F* ratio by dividing the mean using the following equation:

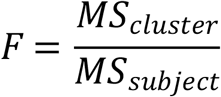

To assess statistical significance, Winer [50] recommended establishing a relatively high α criterion (i.e., 0.20 to 0.30). Generally, however, and in the absence of calculating and testing model ICCs to suggest otherwise, the nested data structure should be modeled using an appropriate analytic technique.

Our study considered the utility of a random effects ANOVA to appropriately account for nested data. Cluster means, an approach historically recommended for handling nested data in preclinical research [e.g., 21–22], however, merit further consideration. Cluster means are an inherently simple approach by which multiple observations within a cluster are reduced to a single, independent observation via the calculation of a summary statistic (e.g., mean; [27, 51]). The validity of cluster means is evidenced by their ability to effectively reduce the probability of type 1 error [18, 27–28]; albeit further research is needed to assess their utility in studies with more complex statistical analyses (i.e., ANOVA). However, when both the number of level-2 units and effect size is small [28], researchers should be cautious about implementing cluster means, as this approach may decrease statistical power.

Generalized estimating equations [GEE; 52] offer another analytic approach for multilevel data. In GEE, statistical corrections are utilized to produce standard error estimates via a ‘sandwich’ estimator, and in some cases parameter estimates, that account for the nested experimental design [52–53]. Unlike ANOVA techniques, GEE are appropriate for non-normal, binary, and categorical dependent variables. When the number of level-2 units is large, compelling evidence for unbiased parameter and standard error estimates supports the validity of GEE for the analysis of clustered data [e.g., 54–56]. However, when the number of level-2 units is small, as is commonly seen in preclinical studies, GEE are too liberal (i.e., increased type-1 error rates, negatively biased standard errors; [e.g., 27, 56–57]). Furthermore, GEE are strictly a population-level modeling approach, which precludes cluster-specific inferences. Thus, although GEE afford a valid approach for modeling multilevel data, they may be impractical for preclinical studies.

Methodological advancements and widely available statistical software packages (e.g., SAT/STAT Software 9.4; SPSS Statistics 26, IBM Corp.) have made appropriately modeling multilevel data readily accessible. Fig 6 offers a recommendation for determining an appropriate statistical approach for the analysis of multilevel data in preclinical studies. Specifically, researchers should begin by calculating ICC and conducting a preliminary statistical test evaluated against a relatively high α criterion (i.e., 0.20 to 0.30; [50–51]). If the ICC is not statistically significant, and the number of level-1 units per cell is small, scientists may conduct a fixed effects ANOVA. However, if the ICC is statistically significant, we recommend accounting for the nested data structure using an appropriate analytic technique (e.g., random effects ANOVA, cluster means, GEE) and any necessary bias corrections (i.e., GEE with small-sample data; [58–59]).

**Fig 6.**
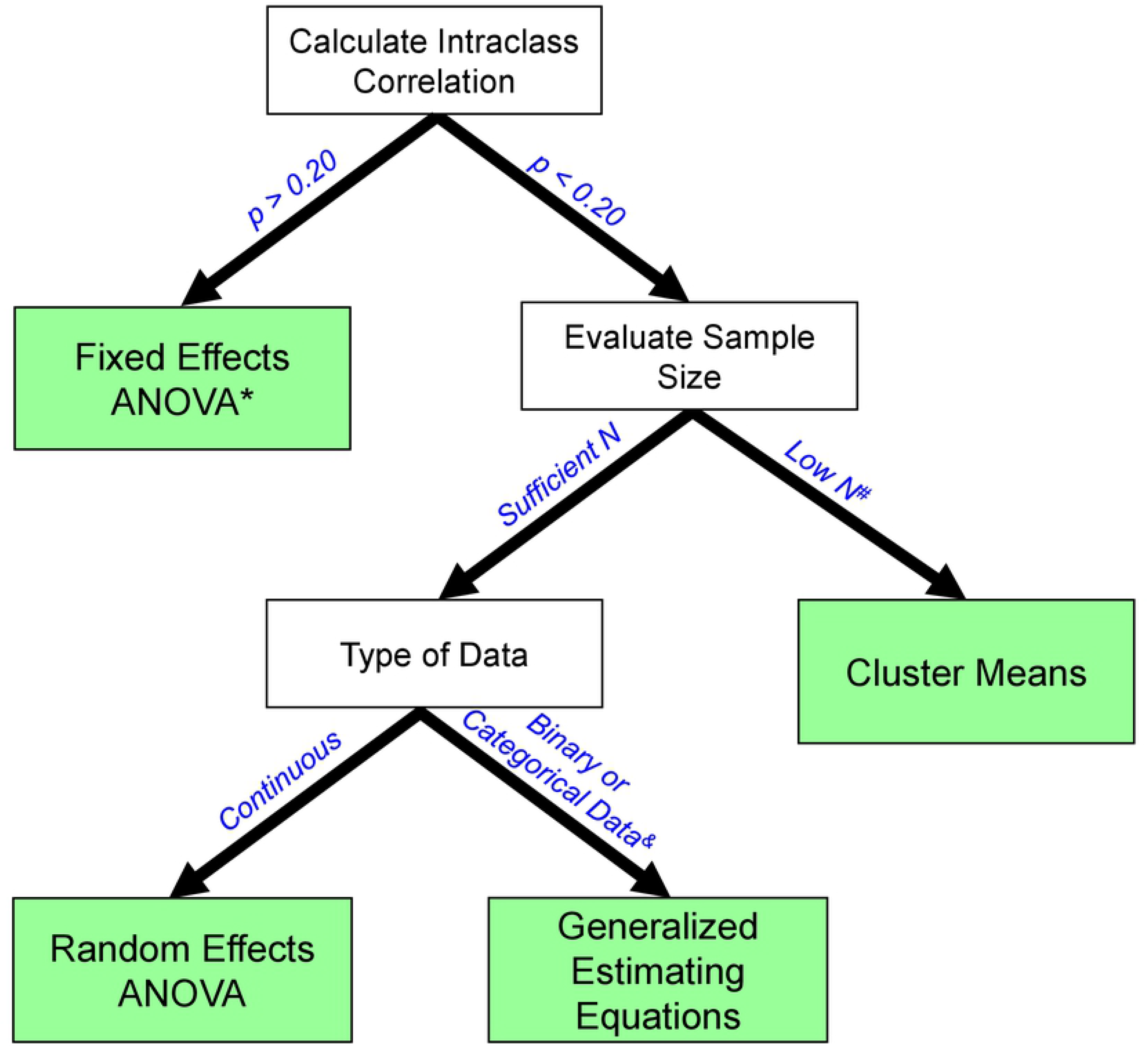
Recommendations for the selection of an appropriate analytic technique for clustered data. A statistical decision tree illustrates some of the key considerations for determining the most appropriate statistical technique for nested data. Critically, these recommendations are not exhaustive, and other statistical approaches may be appropriate dependent upon the research question. *To conduct a fixed effects ANOVA, you will also want the number of level-1 units per cell to be small. #Low *N* in the presence of a large intraclass correlation likely indicates low statistical power. &For preclinical studies with small samples, bias corrections [58–59] may be necessary.

Taken together, the present simulation empirically demonstrates how the failure to account for a nested experimental design may threaten reproducibility in preclinical science. Appropriately accounting for multilevel data via a random effects ANOVA, however, improved the accuracy of both hypothesis testing and parameter estimates. Valid analytic strategies have been provided for a variety of design scenarios to aid in the selection of appropriate statistical techniques for clustered data. Given the prevalence of clustered data in preclinical studies, increased awareness of the implications of inappropriately analyses will lead to enhanced efficiency and translatability.

## Methods

### Experimental Design

#### Population model

The population model in the simulation was a fully crossed 2×2 random effects ANOVA model, with two binary predictors and an interaction term. The level-1 random-coefficients model was defined as follows:

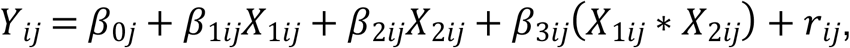

where *β*_*0,j*_ is the intercept, β_1*ij*_ is a level-1 predictor (e.g., Treatment) relating *X*_1*ij*_ to *Y*_*ij*_, β_2*ij*_ is the regression coefficient relating *X*_2*ij*_, a second level-1 predictor (e.g., Biological Sex), to *Y*_*ij*_, β_3*ij*_ is the regression coefficient relating the interaction of the two level-1 predictors (*X*_1*ij*_ * *X*_2*ij*_) to *Y*_*ij*_ and *r_ij_* is the level-1 random effects.

All level-1 coefficients were permitted to randomly vary, yielding the following unconditional level-2 random-coefficient model equations:

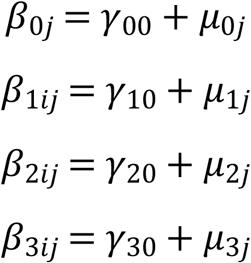

where *γ*_00_ is the average intercept across clusters and *γ*_10_, *γ*_20_, and *γ*_30_ are the average regression slopes across those clusters, corresponding to each given predictor in level-1, respectively, and *μ*_0*j*_, *μ*_1*j*_, *μ*_2*j*_, and *μ*_3*j*_were the associated error terms for each equation.

#### Data Generation

Data for the binary predictors were generated based on a balanced cells design with an effects coding scheme of −.5 and .5 to center the variables. The level-1 coefficients were generated from a multivariate normal distribution using the MASS package and mvrnorm function in R [60]. The mean structure (i.e., fixed effects) was manipulated according to different sizes of the coefficients. The covariances of the level-2 error terms were set to be zero.

The level 1 error term was generated from a normal distribution with a homogeneous variance across clusters (i.e., *r_ij_* ~ N(0, σ^2^)). Variances for both level-1 and level-2 error terms were manipulated to yield the target levels of ICC. The R Foundation for Statistical Computing (version 3.4.1, Vienna, Austria) was utilized to conduct the statistical simulation. The detailed simulation conditions are summarized below.

#### Simulation conditions

Simulation conditions were selected to reflect varying level-1 sample sizes (*N*) and level-2 cluster sizes (*C*) commonly observed in preclinical studies [41–42]. The population value for the model intercept was set to zero. To investigate the impact of variably sized treatment effects, as well as varying size of the interaction between treatment effects and biological sex, parameter values for β_1_ and β_3_ were systematically varied as follows: Null (0), small (0.14), medium (0.39), and large (0.59) [32]. The parameter value of β_2_ was constrained to be 0.14, to focus investigation on detecting variably sized treatment effects of the primary predictor and the interaction term.

Levels of ICC were manipulated by altering the variances of both level 1 and level 2 error terms. Two levels of ICCs were considered, including a small (0.16) and large (0.60) cluster effect. The ICCs were based on the unconditional model. It is noted that the ICC for a given condition may not be identical to the target values. For the small cluster effect, the population ICCs ranged from 0.152 to 0.166 across conditions. In terms of the large cluster effect, the population ICCs ranged from 0.590 to 0.604. Detailed information regarding the population values of the error variances and ICCs is provided in the supplementary materials.

### Statistical Analysis

The nlme: Linear and Nonlinear Mixed Effects Models package [61] in R was used to estimate the random effects ANOVA model. The fixed effects ANOVA model was estimated using the glm function within the same package. A five-way 6 x 5 x 4 x 4 x 2 ANOVA was implemented for post-hoc analyses to analyze the influence of each parameter, and all possible interactions among the parameters, on outcome variables in the study. Given the extremely large sample size, and corresponding inflation of statistical significance, partial η^2^ was utilized to evaluate the practical significance of effects in the study. Specifically, practical significance was evaluated against a partial η^*2*^ ≥ 0.01 criterion, indicating that at least 1% of the variance in a given outcome was attributable to the effect of interest [32]. Post-hoc statistical analyses were conducted using SAS (SAT/STAT Software 9.4, SAS Institute, Inc., Cary, NC, USA). Regression analyses were conducted using GraphPad Prism 5 (GraphPad Software, Inc., La Jolla, CA, USA). Figures were created using GraphPad Prism 5 (GraphPad Software, Inc., La Jolla, CA, USA).

## Code Accessibility

All code utilized for the Monte Carlo Simulation is available upon request.

